# Multimodal fusion of brain signals for robust prediction of psychosis transition

**DOI:** 10.1101/2023.10.25.563602

**Authors:** Jenna M. Reinen, Pablo Polosecki, Eduardo Castro, Cheryl M. Corcoran, Guillermo Cecchi, Tiziano Colibazzi

## Abstract

Psychosis symptoms are often evident before diagnosis, suggesting the underlying biology of high-risk status may predict later disease outcomes. However, a single predictor remains unknown, indicating a need for algorithms that integrate complex information. Here, to identify risk and psychosis conversion, we implemented multiple kernel learning (MKL), a multimodal machine learning approach allowing patterns from each modality to inform each other. Baseline multimodal scans (n=74, 11 converters) included structural, resting-state functional imaging, and diffusion-weighted data. Multimodal MKL outperformed unimodal models (AUC=0.73 vs. 0.66 in predicting conversion). Moreover, patterns learned by MKL were robust to training set variations, suggesting it can identify cross-modality redundancies and synergies to stabilize the predictive pattern. We identified many predictors consistent with the literature, including frontal cortices, cingulate, thalamus, and striatum. This highlights the advantage of methods that leverage the complex pathophysiology of psychosis.

## Introduction

Psychotic illness is associated with significant implications for long-term function. Diagnosis often occurs in adolescence or early adulthood, and is preceded by a period of risk, defined by present but sub-diagnostic symptom levels. To improve treatment and understand the mechanisms of psychosis, a substantial body of literature has characterized this at-risk period (“clinical high risk” or CHR)^1^. A diverse set of clinical and biological (e.g., genetic) methods has been used to determine which individuals develop psychosis^2^. In particular, hope has been placed in neuroimaging to identify biomarkers of risk and conversion, which has revealed an extensive list of biomarker candidates. For example, the onset and early stages of psychotic disorders have been associated with alterations in structural brain features, including decreased whole brain gray matter^3^ and hippocampal volume^4^, ventricular enlargement^5,6^, and insular and prefrontal cortex (PFC) abnormalities ^7,8^. Changes in prefrontal and temporal cortical volumes have been associated with faster rates of decline in converters^3,9,10^. Discoveries in diffusion-weighted imaging (DWI) have implicated white matter tracts as markers of high risk status and conversion in the cerebello-thalamo-cortical circuit^11^ and superior longitudinal fasciculus^12,13^. Functional magnetic resonance imaging (fMRI) studies have shown abnormal PFC-amygdala connectivity in CHR groups^14,15^, alterations in dorsolateral prefrontal cortex associated with working memory deficits^16^, and in functional connectivity (FC) in the cortico-striatal and thalamo-cortical regions that predict increased risk and disease onset^17–20^. Finally, hyperactive cerebello-thalamo-cortical circuitry may predict time to diagnosis^21^.

These numerous predictors vary across modalities, representing only a portion of features associated with risk and conversion. A reliable single-modality, individual-level marker of diagnosis remains elusive, likely reflecting the heterogeneity of psychosis. To address this complexity, machine learning (ML) algorithms have been increasingly used over the last decade to obtain a formal prediction of conversion^22,23^. ML has an advantage in handling complex data using pattern recognition and multivariate analyses, has demonstrated improved performance relative to traditional statistical approaches in predicting conversion^1,24^ and in classifying conversion using neuroanatomical^10,25–27^ and functional imaging features^28^. Promising evidence has shown that adding modalities improves classification in psychosis^23^. Though the number of publications using multiple modalities to predict outcomes is increasing^29^, approaches vary widely. Data fusion methods, which make predictions by combining modalities, are relatively underutilized in brain imaging^30^. For some methods, multimodal information boosts predictive power by optimizing the combination of each modality’s contribution to a final prediction, thereby reducing noise-related uncertainty, as in voting schemes (late fusion)^29,31^. But predictive patterns from different modalities can inform each other during training, allowing complementary information and synergies to produce more robust patterns for each modality. Its simplest implementation is directly concatenating all features (early fusion), treating the whole as one large modality. This can become problematic when collapsing across datasets with unbalanced dimensionality (e.g., neurocognitive assessment vs. whole-brain FC)^32^ where low-dimensional features may be underrepresented when juxtaposed with high-dimensional ones. One solution is intermediate fusion, such as multiple kernel learning (MKL), a simple approach that utilizes a similarity measure (kernel) for each modality across a sample set during training. This makes the dimensionality of individual modalities less relevant yet still allows them to inform each other during learning for increased robustness and performance.^33–35^ Consequently, we expected MKL would perform equivalently or better than single modalities or early fusion when predicting outcomes from multimodal data in schizophrenia. To address this, we used structural, resting-state fMRI, and diffusion neuroimaging modalities within a cohort of healthy participants and CHR individuals followed for two years. We assessed classification in: 1) healthy vs. all CHR individuals; and 2) CHR-converters vs. all individuals who did not develop psychosis (healthy controls + CHR-nonconverters). We compared methodologies by providing prediction accuracy of a support vector machine (SVM) within each modality and two multimodal approaches: 1) direct concatenation of features (early fusion) with SVM; and 2) MKL to compare feature sets with highly disparate dimensionalities. We discuss the benefits, limitations, and contributions of these approaches to understanding the etiology of psychosis.

## Methods

### Participants

Participants (initial recruited sample n = 101) included 61 CHR individuals, of whom 14 were known to convert to psychosis within two years, and 40 healthy individuals similar in demographics who were drawn from the same source population (Table 1). CHR individuals were participants in a clinical research program, the Center of Prevention and Evaluation at the New York State Psychiatric Institute. 74 individuals (42 CHR, including 11 CHR-converters and 32 healthy individuals) had available data from all imaging modalities that passed quality thresholds. CHR status was defined using the Structured Interview for Psychosis-Risk Syndromes /Scale of Prodromal Symptoms (SIPS/SOPS); distinct analyses and additional details from this cohort are reported elsewhere ^15,20^. Among the 42 CHR, at baseline, 14 were prescribed psychiatric medication, of whom 9 were prescribed antipsychotic medication. CHR participants were followed to assess clinical outcomes for 2.5 years. All procedures were approved by the New York State Psychiatric Institute IRB and informed consent was acquired prior to participation.

**Table 1.**
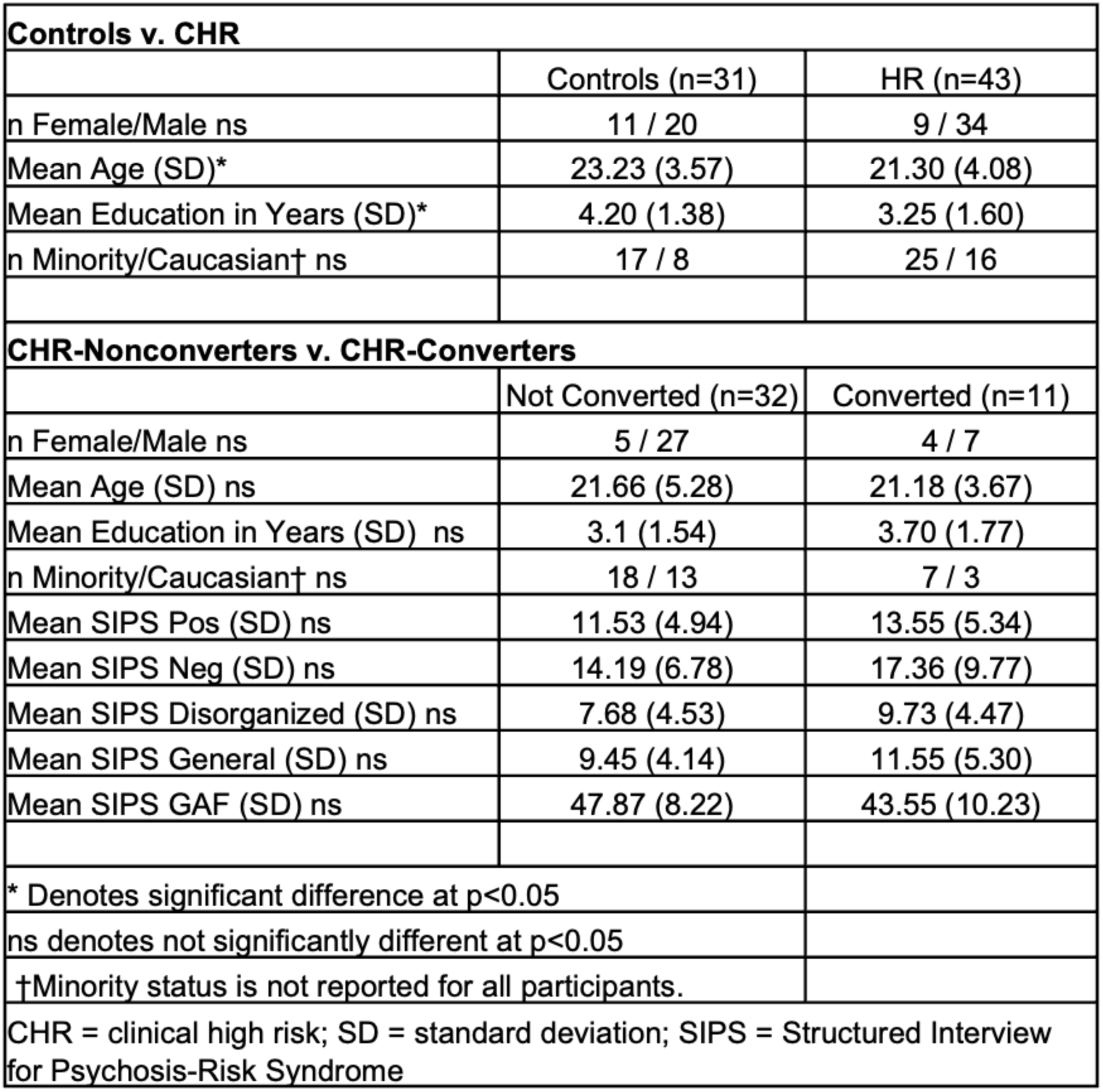
Demographic and symptoms across groups. . Mean and standard deviation (SD) for demographics and symptoms are reported. Chi-square tests were used to assess gender and minority status. T-tests assessed all other comparisons.

### Imaging Acquisition and Preprocessing

Structural, functional (fMRI), and diffusion-weighted (DWI) imaging data were acquired using a GE Signa 3T whole-body scanner using a GE quadrature head coil. Imaging acquisition parameters are described elsewhere ^20^ and in the Supplementary Materials. FMRI and DWI imaging preprocessing was performed using an in-house pipeline created with Nipype ^36^ using Freesurfer v5.3 ^37^ and FSL ^38^. Further detail on this preprocessing, as well as information about the construction of functional connectivity (fMRI FC), fractional anisotropy (DWI FA) and mean diffusivity (DWI MD) summary features are described in the Supplementary Materials.

### Additional Quality Control and Cohort Information

Functional data was first evaluated for quality, and features were then created from qualifying data using partial calculations. In the full sample of 101 participants, 19 had missing functional or structural files, and 8 had files that did not pass quality control thresholds as administered by standard fMRIPrep processes ^39^. Following preprocessing, data were calculated across each resting state scan using a map of 100 ROIs ^40^.

Volumetric structural features were calculated for cortical and subcortical regions using FreeSurfer’s automated cortical surface reconstruction and subcortical segmentation of neuroanatomical regions ^37^. Roughly, this processing incorporates motion correction, removal of non-brain tissue, segmentation of subcortical white matter and deep gray matter structures, and cortical segmentation. Following this process, scans underwent an automatic quality assessment using FreeSurfer’s QA tools ^41^. Volumes that were not adequately processed based on subcortical reconstruction processes or that did not surpass a signal-to-noise (SNR) value of 16 were excluded from our analysis.

### Predictive Models

Our goal was to predict and identify markers of CHR and psychosis conversion. To this end, we focused on two models of interest (Figure 1), classifying: 1) the high-risk label overall (CHR-nonconverters + CHR-converters) versus healthy controls, and 2) theCHR-converter label versus CHR-nonconverters. An analysis examining the converter label (CHR-converters) versus all others (controls + CHR-nonconverters) is reported in the Supplementary Material. In Step 1, a unimodal analysis (Figure 1, top-right) was performed to determine the presence or absence of predictive information for each modality (structure, fMRI, DWI). Here, SVM was used to assess the performance of each modality in both models. The second analysis tested two types of multimodal models (Figure 1, bottom-right). Step 2 (Figure 2, bottom) used a collapsed representation with SVM for a direct concatenation of features, also known as early fusion, to assess prediction for each imaging modality. Finally, Step 3 consisted of a multiple kernel learning model ^33^ for multimodal fusion using EasyMKL ^34,42^. This approach learns a kernel from each individual modality and an optimal combination of them simultaneously. To assess performance, we used a Monte Carlo cross validation scheme with 100 random splits into train and test sets (75%/25%, respectively), using the same proportion of labels in each set. We computed the area under the curve (AUC) of the receiver operating characteristic (ROC) curve and report the average AUC across splits. This was compared to a baseline defined as the same analyses using a random permutation of the test labels. To visualize the variability of learned patterns of each model, we extracted the weight vectors for each modality in each fold, normalized each vector to unit norm, and used principal component analysis (PCA) to project the vectors to two dimensions. We report top feature weights based as those feature labels (brain regions or features) that were given the highest absolute weights, on average across folds.

**Figure 1.**
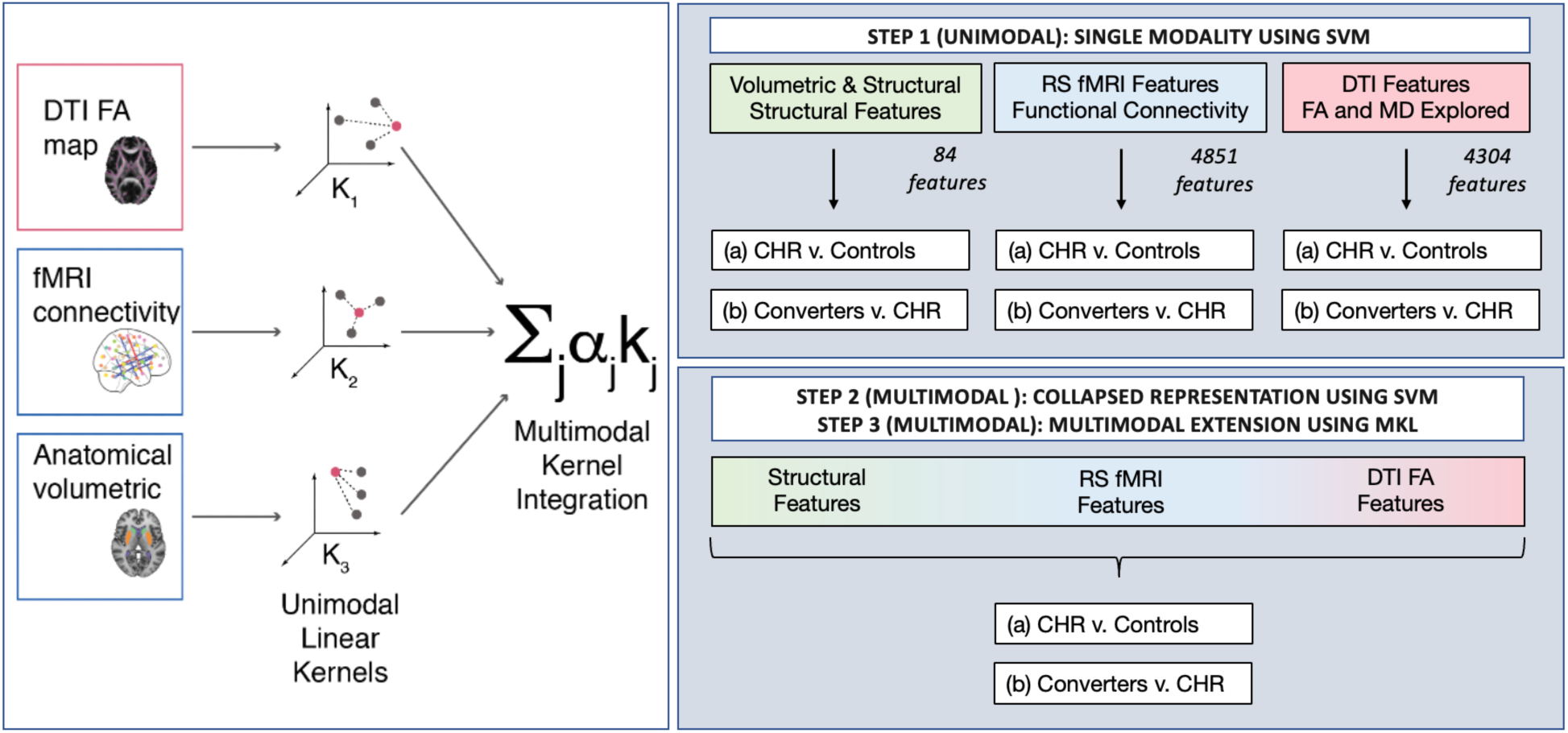
Analysis Specifications and Flowchart. (Left) Multiple kernel learning (MKL) is a part of the family of kernel methods. These make a prediction about a test sample based on its similarity (kernel function) to samples seen during training. In multimodal multiple kernel learning, schematized here, a multimodal kernel is learned as a combination of kernels obtained from single modalities, allowing patterns to be learned from each modality to inform the rest. (Right) Predictive models were assessed with a unimodal analysis where each data modality was evaluated individually to predict the given label. Next, multimodal analyses are shown using support vector machines (SVM), and finally, a MKL model. *RS=resting state; DTI=diffusion tensor imaging; FA=fractional anisotropy, MD=mean diffusivity; CHR=clinical high risk*.

**Figure 2.**
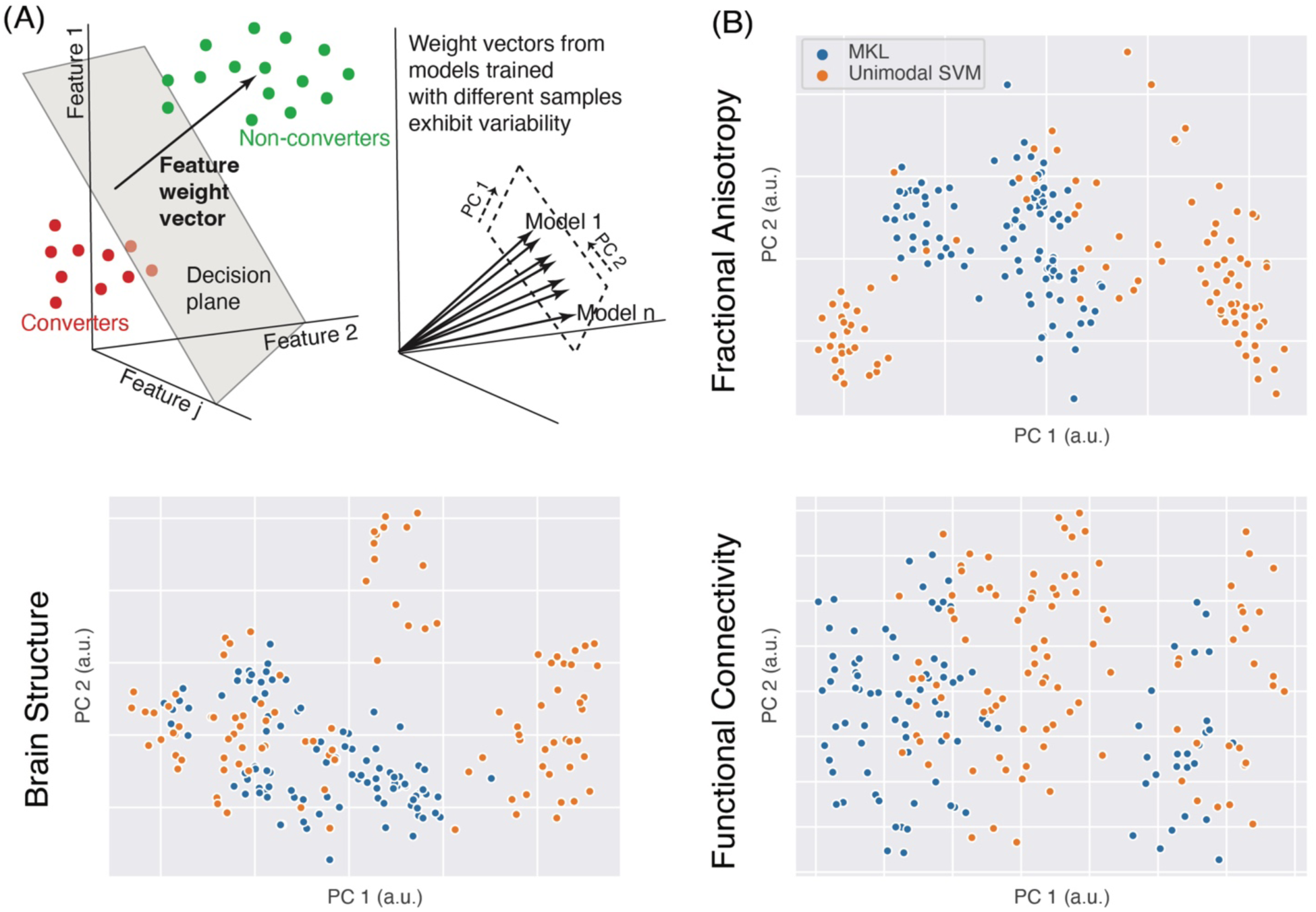
Variability of brain patterns learned by unimodal and multimodal models. (A) Linear classifiers use feature weight vectors for making a classification, representing the multivariate patterns of feature importance and variability across training samples and folds. (B-D) We used principal component analysis (PCA) to visualize this weight vector variability across training folds for each modality in unimodal SVM and MKL when predicting conversion. Relative to SVM models which formed clusters based on training samples, MKL benefitted from other modalities to stabilize the feature weights.

## Results

### Participant Characteristics

We first addressed demographic and motion characteristics within and across the groups. During participant recruitment, proportional matching was performed for race, sex, and age of the CHR and healthy individuals. In the present analysis, we specifically examined subgroups of the original cohort based on data availability, quality, and specific comparison (e.g., CHR or healthy individuals from the original cohort who had data from all modalities that passed QC). As a result, we re-assessed demographics, and where appropriate, symptoms (see Table 1) for each cohort comparison. A comparison across healthy controls and CHR participants showed that there were differences in age and education (both p<0.05), though no differences were identified for CHR-converters versus CHR-nonconverters for demographics or symptoms. Because symptoms (SIPS/SOPS), age, and sex were similar across CHR-converters and CHR-nonconverters in this cohort (Table 1), they were not used as variables of interest. Instead, we focused on brain measures alone. No differences were found between the groups for motion related to the BOLD signal; head motion defined by mean displacement was not different between healthy control and CHR (t(71)=-0.62, p=0.54), or CHR-nonconverters and CHR-converters (t(40)=-1.86, p=0.07).

### Comparison of Model Performance and Stability

We then examined the performance results of machine learning classifications, first for the CHR label (CHR-converters and CHR-nonconverters) versus the control label. FA and fMRI FC features had the best performance of the single feature modalities (Table 2; AUC=0.61, p=0.027 and 0.62, p=0.02, respectively). MD and structural features were not significant predictors. The MKL model tied with early fusion SVM in this instance with AUC=0.66, p<0.01. Next we examined the analysis classifying the CHR-converter versus CHR-nonconverter label. Here, FA and fMRI FC features also yielded the best single-modality predictions (AUC=0.66, p=0.043 and AUC=0.6, p=0.037, respectively). Similar to the first analysis, MD and structural features were not significant. The MKL model outperformed any single modality and early fusion SVM with AUC of 0.73, p<0.01. This suggests a performance advantage for multimodal models, and highlights the importance of FA and fMRI FC in predicting psychosis onset.

**Table 2.**
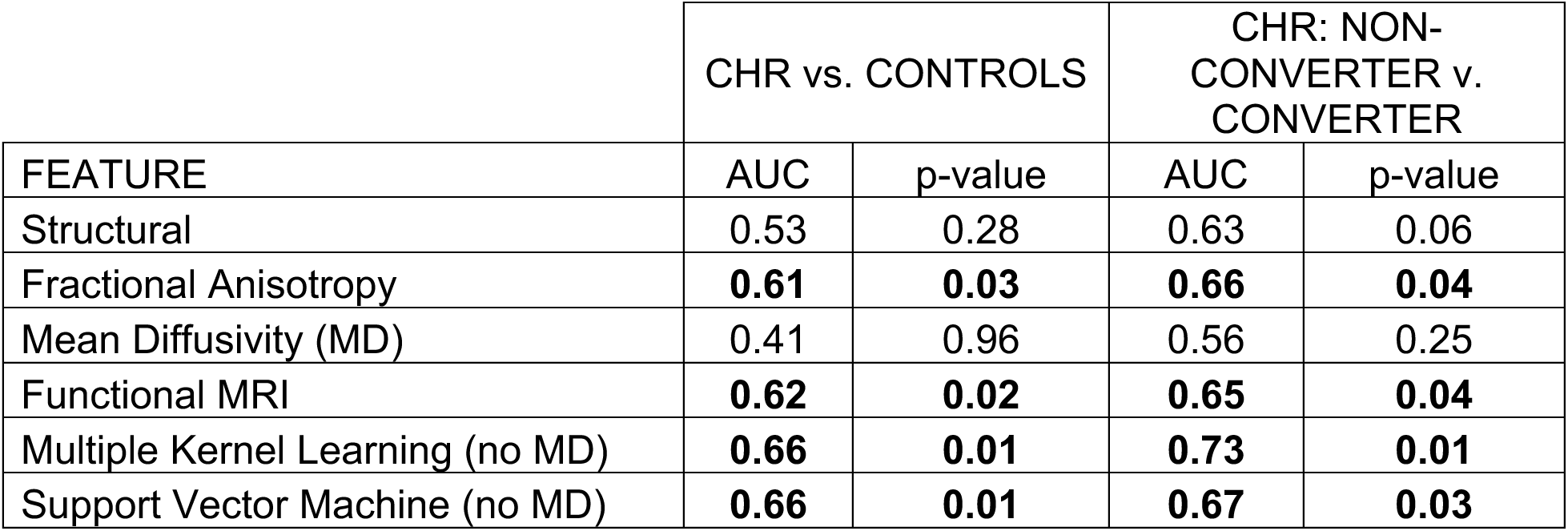
Model performance. . Model performance was quantified for each analysis. The MKL model demonstrates superior performance when identifying converters. *Bolded values denote significance at p<0.05. CHR = Clinical High Risk*

| FEATURE | CHR vs. CONTROLS |  | CHR: NON-CONVERTER v. CONVERTER |  |
| --- | --- | --- | --- | --- |
|  | AUC | p-value | AUC | p-value |
| Structural | 0.53 | 0.28 | 0.63 | 0.06 |
| Fractional Anisotropy | **0.61** | **0.03** | **0.66** | **0.04** |
| Mean Diffusivity (MD) | 0.41 | 0.96 | 0.56 | 0.25 |
| Functional MRI | **0.62** | **0.02** | **0.65** | **0.04** |
| Multiple Kernel Learning (no MD) | **0.66** | **0.01** | **0.73** | **0.01** |
| Support Vector Machine (no MD) | **0.66** | **0.01** | **0.67** | **0.03** |

To understand the effects of multimodal fusion in learned patterns and their stability, we assessed their variability in the unimodal (SVM) and multimodal (MKL) models. We considered how the weight vectors (i.e. the vectors quantifying the signed importance given to each feature in a given modality, Figure 2A) changed when different subsets of the data were used on each training fold, and visualized how they were distributed using principal component analysis to project them to two dimensions (Figure 2 and Supplementary Figure S9). Unimodal SVM models tended to be multi-stable, forming clusters depending on the samples seen during training. By contrast, MKL weights tended to benefit from the presence of other modalities, which stabilized the feature weights (especially FA and structural maps). A similar behavior was observed in CHR prediction (Supplementary Figure S9). This suggests that multimodality can ameliorate the stability issues of high-dimensional patterns that are typical of the low-sample-number brain models common in psychiatric disorders.

### Features Predicting Healthy controls vs. all CHR Participants

Next, we examined features predicting the CHR label (CHR-converters and CHR-nonconverters) versus control label using the MKL model (Figure 3, upper panel). The top structural features predicting CHR status for the SVM model included regions of cortex, including postcentral gyrus, cingulate, and insula, as well as enlarged ventricles and decreased dorsal striatum and frontal cortex volume (Figure 3A and Supplementary Table S3). Additionally, reduced fMRI connectivity in limbic, striatal, fronto-cortical and cerebellar regions, including connectivity in medial and inferior frontal gyrus and caudate, also predicted the CHR label, as did heightened thalamic-cortical and fronto-striatal connectivity (Figure 3B; see Supplementary Materials for single-modality findings). The top FA features predicting the CHR label included decreases in corpus callosum and limbic regions (Figure 3C). Differences between MKL-SVM models indicated the MKL model leveraged SVM predictors by identifying the optimal combination of each modality, including reduced volume in frontal and temporal cortex, ventricle size, and fronto-limbic functional connectivity (Supplemental Figure S1).

**Figure 3.**
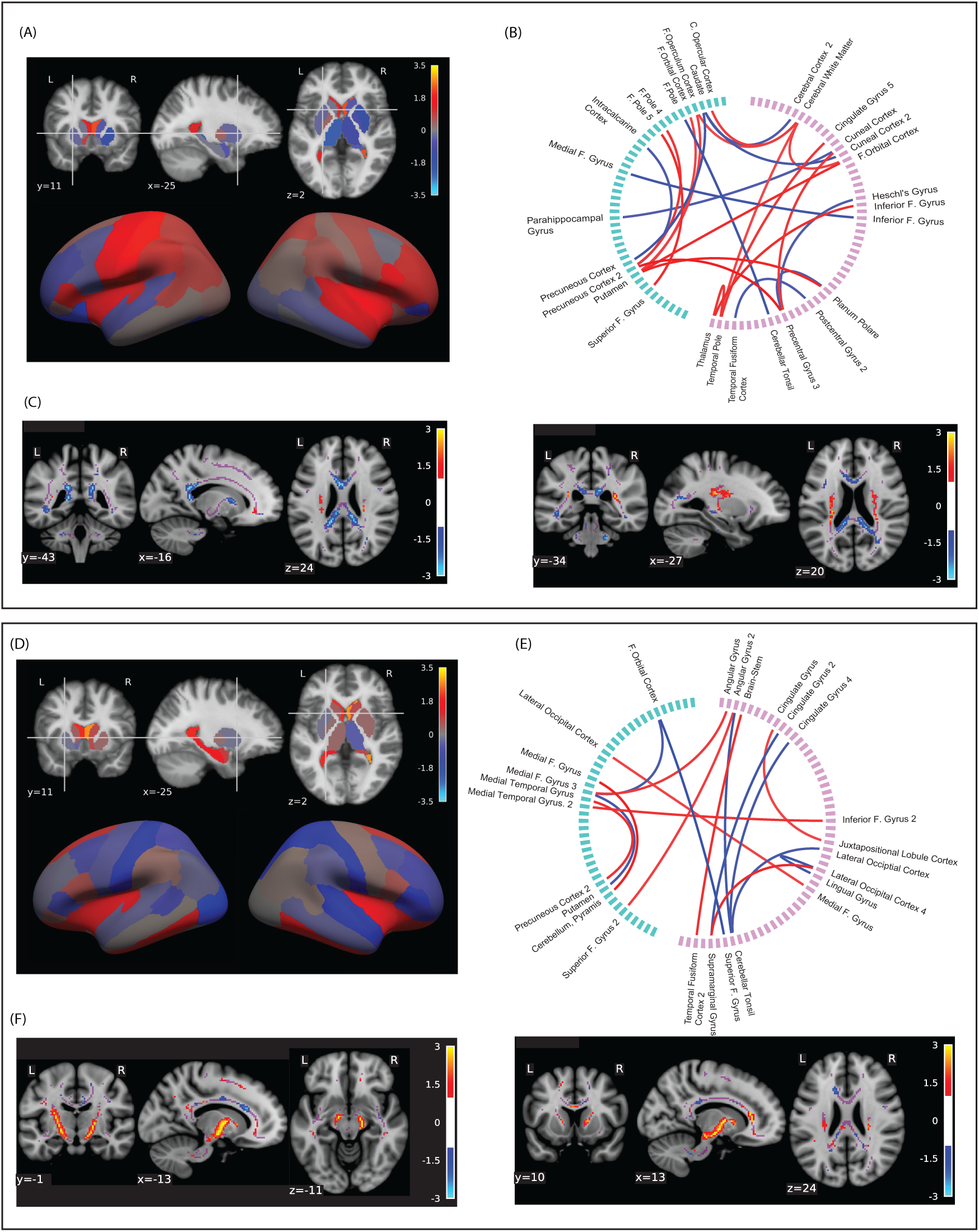
Top features predicting (above) CHR vs. Controls and (below) CHR-Converters vs. CHR-Nonconverters using the MKL model. Top feature weights are shown for (A, D) subcortical and cortical structural features; (B, E) functional connectivity features for which green = L and pink = R, where red denotes positive and blue denotes negative connectivity; and (C, F; bottom two panels) fractional anisotropy features. [F = Frontal; C = Central]

### Features Predicting CHR-Nonconverters vs. CHR-Converters

The MKL model predicting the CHR-converter versus CHR-nonconverter label (Figure 3, lower panel) indicated that structural features identifying converters from high risk non-converters included enlarged ventricle size, as well as orbitofrontal, insula, cingulate, and hippocampal volumes (Figure 3D, Supplemental Table S1). Functional connectivity features predicting conversion included decreased connectivity in multiple cortical regions, and between cortical and striatal regions as well. Both reduced and heightened connectivity predictors were also identified in various cortical regions, including frontal, temporal, cerebellar, and cingulate cortices (Figure 3E). When examining DWI predictors of conversion, we identified notable increased bilateral thalamic FA as a top predictor of psychosis, providing further evidence for the role of this region in predicting conversion (Figure 3F). Differences between the MKL-SVM top features indicate that MKL models further leveraged ventricle size, decreased frontal and parietal cortical volume, and numerous frontal, striatal, cerebellar and limbic functional connections to predict the converter label (Supplemental Figure S2).

## Discussion

Using a multimodal brain imaging dataset comprised of functional, structural, and white matter brain imaging features to predict risk and conversion to a psychotic illness, we demonstrated that a multimodal approach achieves superior performance over a unimodal approach, and in particular that MKL was superior relative to single-modality SVM and collapsed feature SVM in predicting conversion to psychosis from high-risk status. Across our two analyses of interest, MKL leveraged the multimodal neurobiology of psychosis to learn stable predictive patterns that enhance accuracy, a key element in small studies using high dimensional data to predict outcomes. These findings suggest that there may be an advantage in predicting psychiatric outcomes from multiple modalities and in optimizing the combination of variables with an intermediate-fusion approach. This approach reveals clear differences and similarities in the features predicting risk and conversion through the inspection of common features across modalities, for instance in the insula, cingulate, striatum, and along thalamo-cortical tracts. Moreover, these findings have implications extending to modalities beyond neuroimaging. For example, MKL could be particularly useful when combining cognitive and genetic features with neuroimaging data, each of which is expected to have distinct feature dimensionality and appropriate similarity measures. Collectively, these findings point to the value of using a multimodal approach, and more specifically, that using MKL with multimodal data to understand and predict risk and conversion offers benefits beyond other methods.

Unlike ML approaches that complicate the ability to inspect learned patterns, one advantage to this method is that it does not obfuscate the brain regions involved. To the contrary, the increased stability of learned patterns in MKL allowed us to explore the top predictive features within the models and assess their consistency with prior findings. For instance, when examining risk alone, we found that frontal, striatal, and thalamic dysconnectivity predicted the CHR label (CHR-nonconverters + CHR-converters) versus controls, converging with growing evidence that these regions serve as biologically meaningful markers of risk and psychosis^18,43^. Structural features predicting CHR status further verified this, as volume of the thalamus, putamen, frontal and cingulate cortex, and sensory and motor cortices were also CHR predictors. Fractional anisotropy features distinguishing CHR from controls included negative weights in the cingulate. Comparing across modalities, the cingulate, frontal, and somato-motor cortices, striatum, and thalamus emerged as multimodal markers of CHR, aligning these findings with prior literature^43,44^. When predicting conversion, many regions were characterized by both hypo- and hyper-FC, including frontal cortical regions, cingulate, and cerebellum, perhaps indicating an overall pattern of dysconnectivity. Structural features predicting conversion included frontal cortex, cingulate, insula, and a range of subcortical regions. Well-aligned with prior converging evidence, ventricle size and hippocampus distinguished converters in the SVM analysis, which was further leveraged in the MKL approach^5^. FA predictors of conversion included robust activation of the bilateral striato-thalamic tract, and negative weights in corpus callosum and cingulate. Together these observations underscore the flexibility of multimodal modeling to tap into the complex neurobiology of psychosis.

There were several commonalities across modalities, including those in dopaminergic pathways, cerebellar-cortical and thalamo-cortical regions, insula, and cingulate. Volumetric predictors in the striatum, medial temporal lobe, orbitofrontal, and occipital cortices aligned with findings that abnormalities in these regions are associated with illness onset and early first-episode schizophrenia^6,45^, possibly implicating the pathophysiology in dopaminergic pathways, a putative mechanism underlying symptoms of schizophrenia^46,47^. Top feature weights predicting conversion implicated both hypo- and hyperconnectivity in cerebellar-cortical regions, partially aligning with findings in cerebellar hyperconnectivity predicting conversion^21^. Insula volume and FA were top predictors of CHR status, with insular volume also distinguishing converters from other CHR patients, which aligned with prior reports of structural and functional abnormalities in high-risk individuals^8,48–50^. Increased thalamo-cortico tract FA was a top predictor of conversion, also verifying broad findings associating abnormal thalamic function with conversion^51–54^. Finally, the cingulate was a predictor of both CHR status and conversion as evidenced by including dysconnectivity in the FC data, structural abnormalities, and reduced FA. Overall, this was largely consistent with mechanisms thought to underlie early psychosis, implicating a diverse set of brain regions and a broad pattern of whole-brain dysconnectivity, affecting a range of functions associated with psychosis. These included primary sensory and motor cortices, regions supporting dopaminergic and salience processing^47^, and frontal regions supporting higher cognitive processes such as working memory and cognitive control^15^. In particular, thalamic abnormalities were multimodal predictors of both risk and conversion, which may reflect compensatory activity aimed to support abnormal salience and sensory processing^54^, resulting in broad downstream physiological changes. This list of regions highlights the complex nature of predicting conversion, and the need for methodologies that account for this complexity in predicting outcome.

The present study also serves as an opportunity to test the accuracy of MKL approach, relative to unimodal models side-by-side. We found that each modality showed different predictive power depending on the classification task, with FA and fMRI performing best when classifying converters. DWI MD-based models did not perform above chance levels in any task and that modality was omitted in multimodal analyses. Generally, we were more successful at distinguishing the converter group than the CHR group. We speculate that this is possibly due to greater heterogeneity in the CHR group, which was recruited with a broad set of clinical requirements, whereas the converter group is ultimately defined by a more narrow set of diagnostic SIPS criteria. Further exploration comparing modalities and performance is needed, as the number of studies using machine learning to predict psychosis onset is small relative to those using traditional statistical approaches. We also advocate here more generally for the use of intermediate fusion methods like MKL. Though studies using multimodal imaging with machine learning to identify schizophrenia or CHR are relatively uncommon^6,22,27,54,55^, and those predicting psychosis onset are even less common^25,56^, many of those published typically used a straightforward SVM approach for single modalities, combined together using a voting scheme (late fusion). We had expected that MKL would boost predictive power by averaging out noise in the prediction of single modalities^29^, allowing learned signal from each modality to inform or affect the others.^33–35^ Here we confirmed that MKL can indeed do that, learning a kernel from each individual modality and generating an optimal combination of them simultaneously. ^57^ It is likely because of it that it performed best in our comparison analyses. This suggests that intermediate fusion methods, of which MKL is arguably the simplest, may have an advantage with multimodal imaging data, which may be even more pronounced in feature sets of heterogeneous size.

This is the first study to use multimodal imaging and MKL to predict psychosis to our knowledge, and there are important considerations regarding possible extensions and limitations. In addition to conversion, future studies should explore variables like symptom remission, functional outcomes, and medication use. Replication in independent datasets should include data with diverse dimensional characteristics, like cognitive, clinical, demographic, and genetic features. In particular, cognitive features may enhance predictive accuracy in psychosis^23^. Here, MKL could show a particular advantage, given the extreme difference in dimensionality between, for example, high-dimensional FC features and scores from lower-dimensional cognitive assessments. Additionally, it could directly accommodate other modalities for which specific measures of similarity are relevant, as with genetic sequences. Finally, though this study had a smaller sample size than others using predictive modeling, we are optimistic that these findings can be replicated, and perhaps accuracies may be amplified with a larger training set.

Overall, we demonstrated clear advantages for the use of multimodal datasets and use of MKL, both in terms of predictive performance and model robustness. Across modalities, we identified many biological features, including the cingulate, thalamus, striatum, and frontal cortices, which emerged as cross-modality predictors of CHR and conversion, providing strong converging evidence for each of these as biomarkers of risk and psychosis onset. This large number of common predictors also highlighted the multidimensionality of mechanisms underlying psychosis, and the strong need for multimodal learning models when predicting outcomes.

## Acknowledgements

This work was supported in part by National Institutes of Health (5U24MH124629 to JMR, PP, ED, CMC, and GC; R01MH107558 and R21MH086125 to CMC).

## Data Availability

The data that support these findings may be available upon request, but are not publicly available and subject to restrictions outlined by the IRB and consent form.

## Author Contributions

JMR: imaging coding and analytics, paper writing, figure creation/editing PP: ML analytics, imaging coding and analytics, paper writing

EC: structural analytics, figure creation, paper writing

CC: oversaw study design and data collection, paper writing GC: oversaw analytics, paper writing

TC: oversaw study design and data collection, paper writing

