## Supplemental Materials for "Multimodal fusion of brain signals for robust prediction of psychosis transition"

### Supplemental Information for: Prediction of high risk status and psychosis transition from robust multimodal brain signals

Jenna M. Reinen*, Pablo Polosecki*, Eduardo Castro, Cheryl Corcoran, Guillermo Cecchi, Tiziano Colibazzi

#### Structural and Functional Imaging Processing

T1-weighted images were acquired using SPGR sequence with the following parameters: TR = 85ms, TE=21ms, 75 degree flip angle, FOV=25cm, receiver bandwidth = 31.2 KHz/sec, 352 x 256 matrix, slice thickness = 5mm. Following that, 3-dimensional spoiled gradient recall image were used for coregistration with the axial echoplanar images (EPI) used for the resting state scans. Resting state EPI were acquired with the following specifications: TR=2200ms, TE=30 ms, 90 degree flip angle, FOV=24cm, receiver bandwidth=250 KHz/sec, single excitation per image, slice thickness=3.5 mm, slices per volume=34; 24 x 24 cm field of view, 64 x 64 matrix, with a resolution of 3.75 x 3.75 x 3.5 mm using whole-brain coverage. For each resting state run (duration: 5 minutes, 21 seconds), participants were instructed to remain still, close their eyes, and allow free mind wandering. Diffusion-weighted MRI images and a baseline (b0) volume were acquired along 25 gradient directions with parameters as follows: b-value = 1000s/mm$^2$, TR = 17000ms, FOV = 24cm, 132 x 128 matrix and slice thickness = 2.5mm.

For functional imaging preprocessing, 5 time points were discarded to account for T1-equilibrium effects. Slice time correction was applied, followed by correction for head motion using rigid realignment to the middle of the time series volume. A threshold of 6 frames of relative displacement (defined using FSL's *mcflirt* above 0.6mm) was a criterion for exclusion, however, this did not affect any of the participants with a complete multimodal dataset. Residual head motion effects and physiological artifacts were corrected using tCompCor [38]. Voxel time series were band-pass filtered (0.01-0.16Hz). Following brain extraction, EPI time series were co-registered to the anatomical volume and nonlinearly normalized to MNI space. Registration was assessed as verified in prior studies [39].

DWI images were preprocessed using in-house Nipype pipelines. Images were corrected for motion, eddy current distortions, and bias [41]. Geometric distortions from magnet artifacts were corrected using nonlinear registration via T1 registration[42]. Images were then fit to a diffusion tensor. Features for individual participants were created by registering each baseline image and computing a TBSS skeleton[43]. Fractional anisotropy (FA) and mean diffusivity (MD) summary features were then projected along the white matter skeleton using tract-based spatial statistics. TBSS maps were downsampled to 2mm isotropic voxels.

**Additional Group-Level Analyses: Comparing Converters to All**

In addition to the analyses described within the main text, we also examined results for an additional analysis that predicted the CHR-converter label relative to all other participants ("converters vs. all"; this is the CHR-converter label compared to all other participants, e.g., including both controls and CHR-nonconverters). These results are presented below and throughout the Supplement, and are compared to the two other analyses (predicting high risk label, and CHR-converters vs. CHR-nonconverters).

When comparing the converter label to the entire cohort of non-converters (including controls and non converter CHR), we found that of the single modalities, the structural features performed best, and it was the only modality that predicted outcome with significance, with AUC = 0.65, p = 0.029. The MKL model was able to outperform the single modalities with AUC = 0.68 and p = 0.01, and MD features (see Supplementary Table S2). The lateral ventricles, regions of frontal cortex, parahippocampal regions, cingulate cortex, and insula were among the top structural predictors using MKL (Supplementary Table S1, Supplementary Figure S3, Panel A). The top functional imaging connectivity features predicting conversion included both heightened and decreased cortical regions, including in frontal, orbitofrontal, and cingulate regions, as well as in putamen and cerebellum (Supplementary Figure Panel 3B). As in the other analysis that predicted the converter label (CHR-converters vs. CHR-nonconverters), increased bilateral thalamic FA was a top feature predicting converters relative to all other participants (Supplementary Figure S3, Panels C&F). When examining the differences between the MKL and SVM models, a comparison revealed that the MKL leveraged similar regions across models, including a similar pattern of structural features, additional frontal and limbic connectivity, and bilateral thalamic FA.

#### Supplementary Figures

#### Additional MKL Analyses

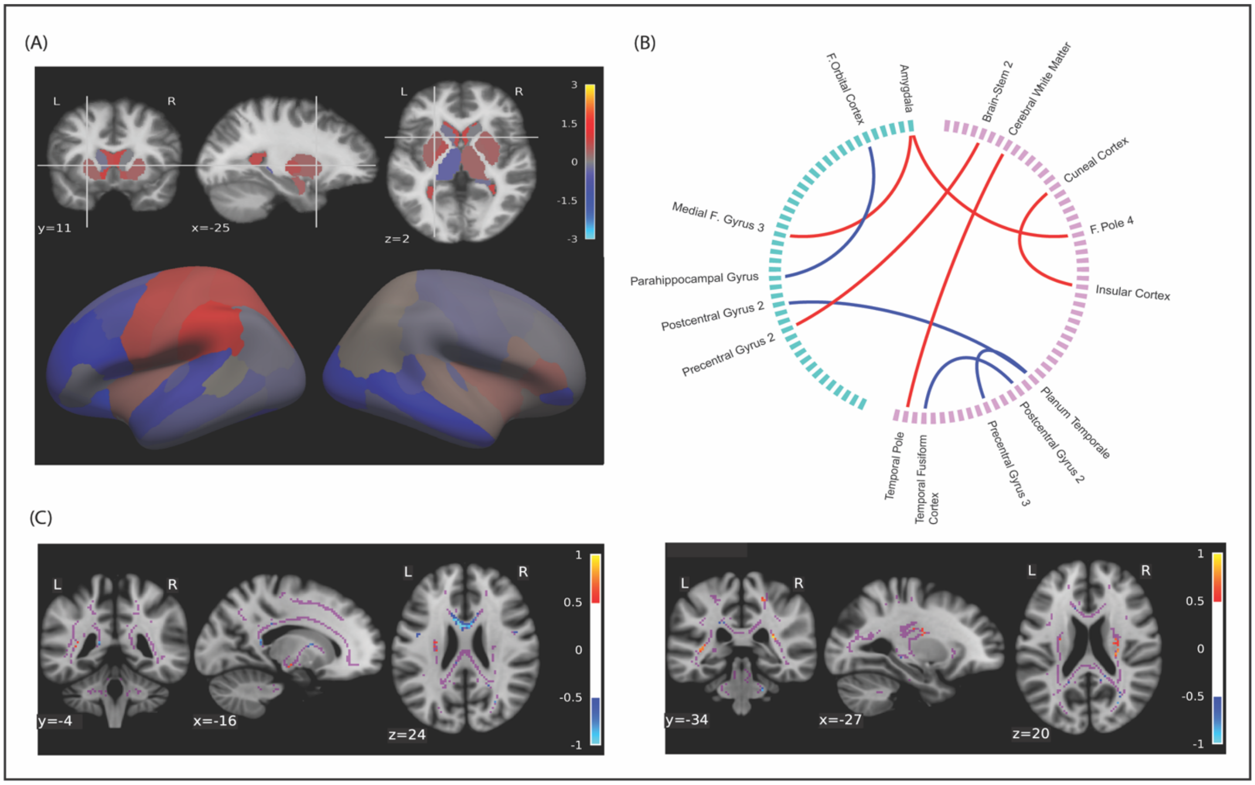

**Supplemental Figure S1. Difference in MKL–SVM Features Predicting the CHR vs. Control label**. A comparison of the in features used in the MKL minus SVM analyses to predict the high risk (CHR-nonconverters + CHR converters) versus control label is shown for (A) structural features; (B) fMRI FC features; and (C) FA features. [F = Frontal.]

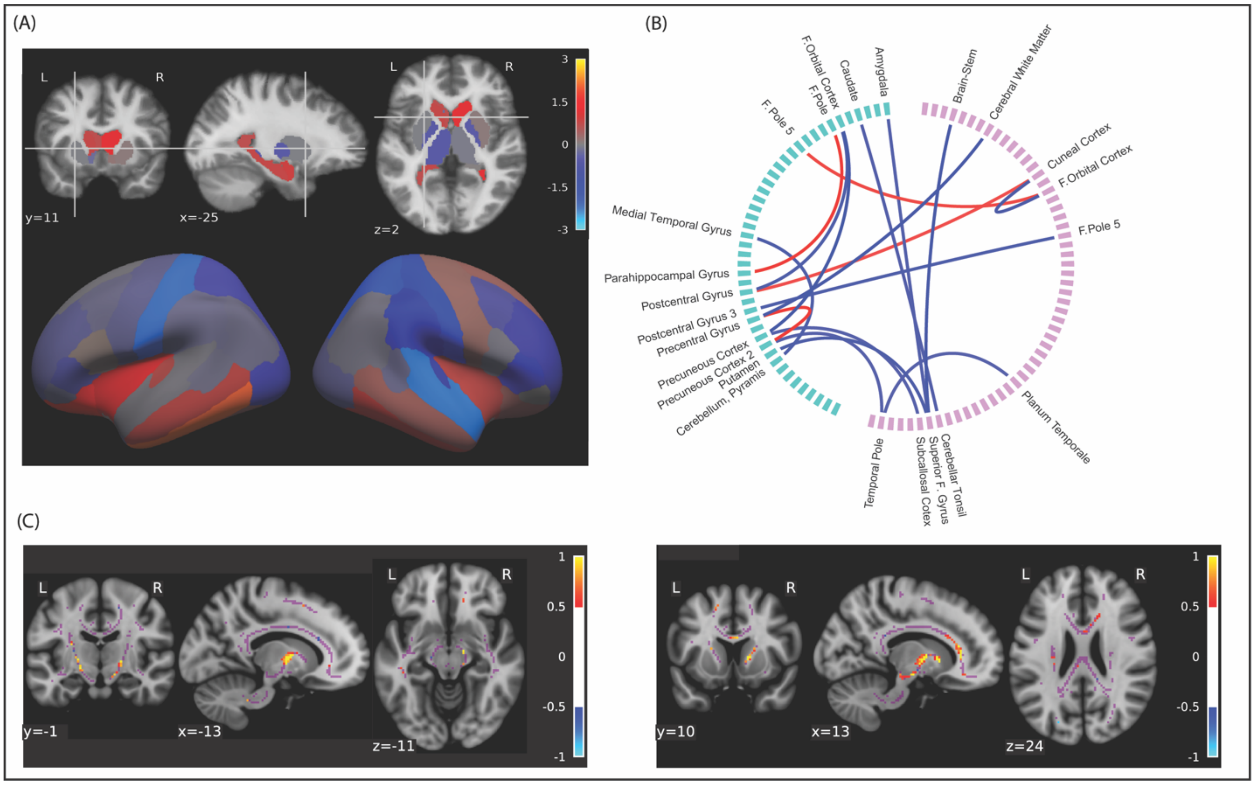

**Supplemental figure S2. Difference in MKL–SVM Features Predicting the Converter Label**. A comparison of the in features used in the MKL minus SVM analyses to predict the converter label (CHR-converters versus CHR-nonconverters) (A) structural features; (B) fMRI FC features; and (C) FA features. [F = Frontal.]

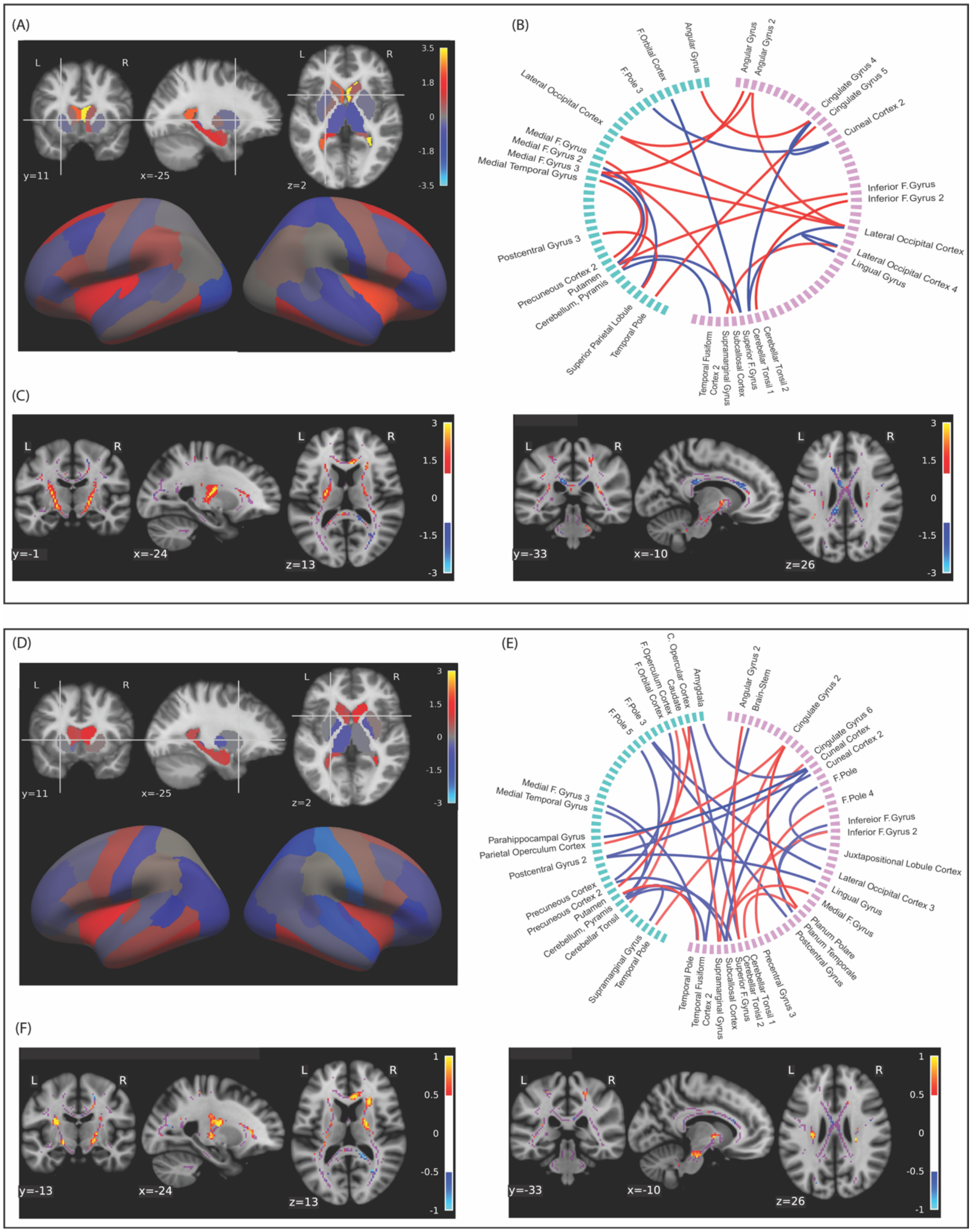

**Supplementary Figure S3. Predicting the Converter Label vs. All Other Participants.** Top feature weights are shown for (A, D) subcortical and cortical structural features; (B, E) functional connectivity features for which green = L and pink = R, where red denotes positive and blue denotes negative connectivity; and (C, F; bottom two panels) fractional anisotropy features. The top panel (A-C) shows results from the MKL model; the bottom panel (D-F) shows the difference between the MKL – SVM models. [F = Frontal]

#### Single Modality Findings

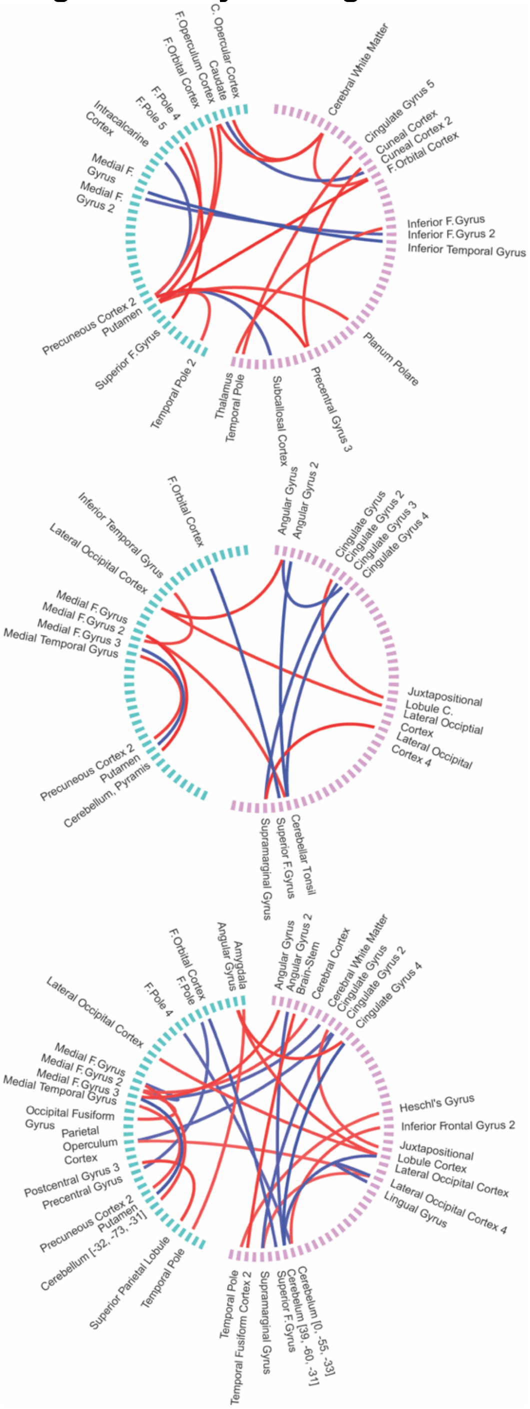

**Supplementary Figure S4. Top Functional Connectivity Predictors from the Single Modality Analyses Using SVM**. Regions of functional connectivity predicting hyperactive (red) connections and hypoactive (blue) connections for the (top) high risk label (CHR-nonconverters and CHR-converters); (middle) CHR-converter label relative to the CHR-nonconverter label; and (bottom) CHR-converter label relative to all nonconverters (controls and CHR-nonconverters). [C = Central; F = Frontal]

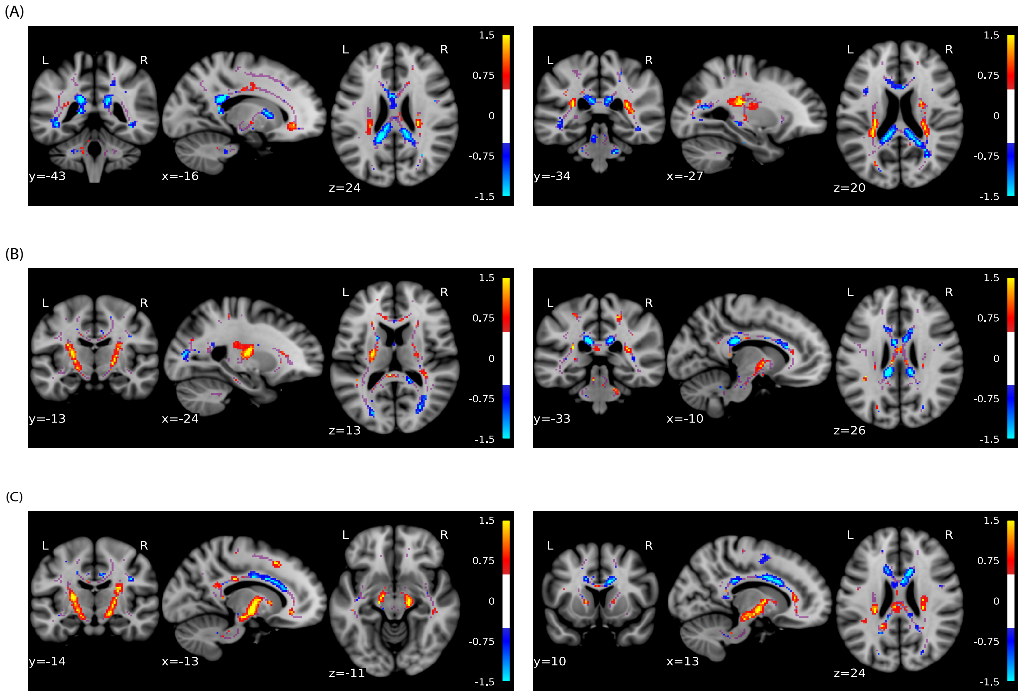

**Supplementary Figure S5. Supplementary Figure 1. Top DWI Fractional Anisotropy Predictors from the Single Modality Analyses Using SVM**. The top feature weights are shown for regions that passed the designated threshold. The plots were derived from the analyses that best predicted (A) the high risk label (CHR-nonconverters and CHR-converters) relative to control participants; (B) the converter label relative to all other participants; and (C) the converter label relative to other CHR participants.

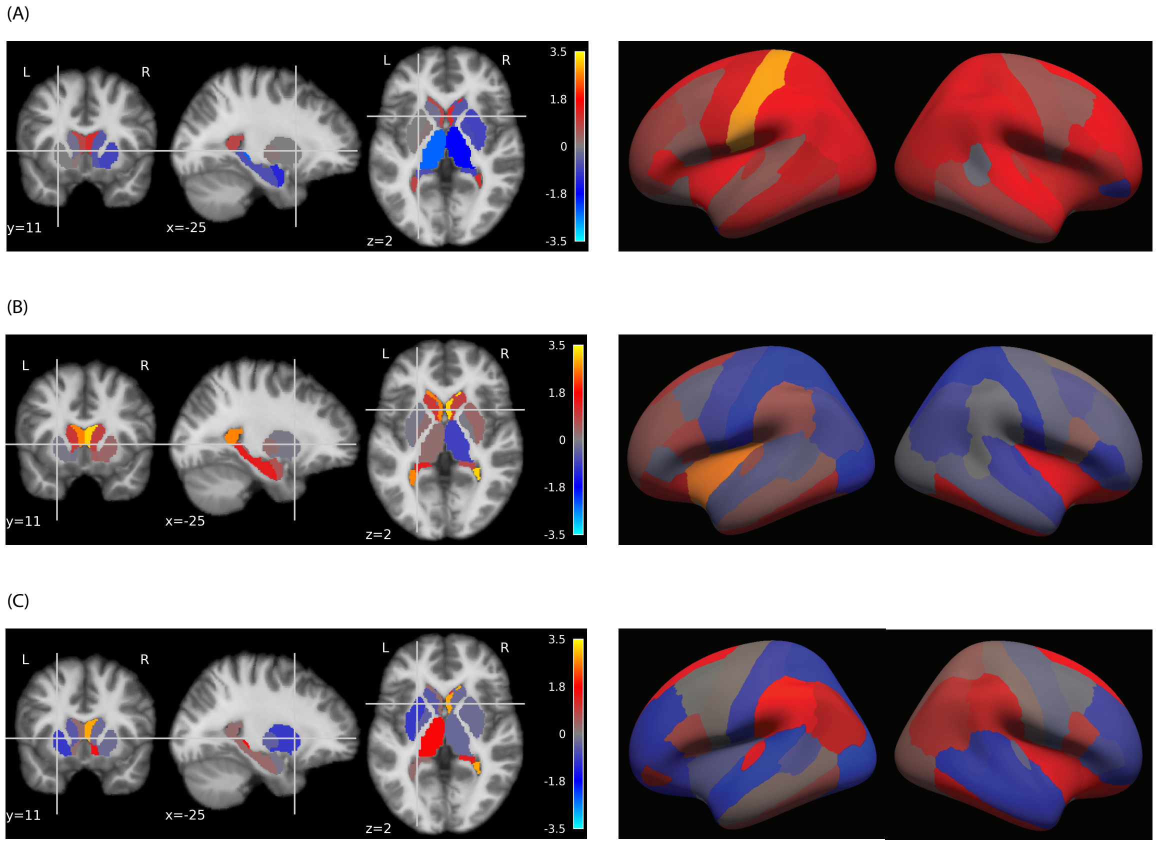

**Supplementary Figure S6. Structural features predicting high risk and conversion to psychosis from unimodal analysis**. Feature weights for structural regions are shown for (left) subcortical structures and ventricles predicting (A) the high risk label relative to control participants; (B) the converter label relative to other CHR participants; and (C) the converter label relative to all other participants. (Right) cortical regions predicting same set of analyses.

#### Section 3. Additional Analyses for the MKL Model

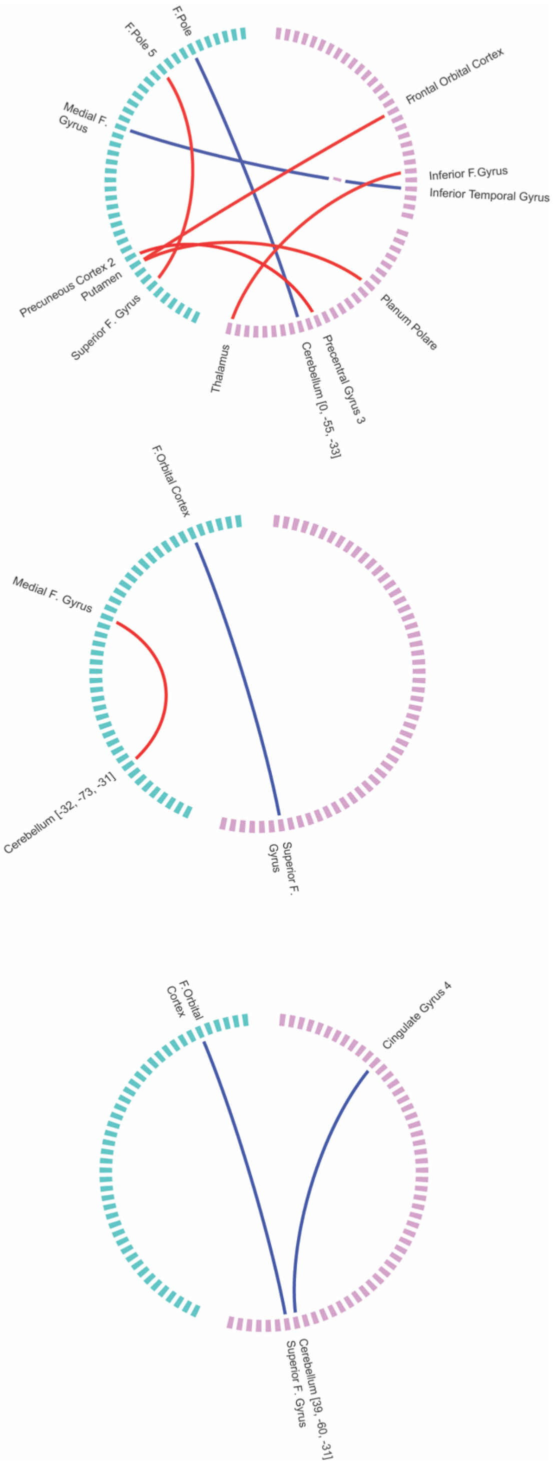

**Supplementary Figure S7. Higher-threshold (conservative) findings in functional connectivity analysis (threshold = 6)** for (top) the high risk label; (middle) the CHR-converter label versus CHR-nonconverter; and (bottom) the CHR-converter label vs all other participants. [F = Frontal]

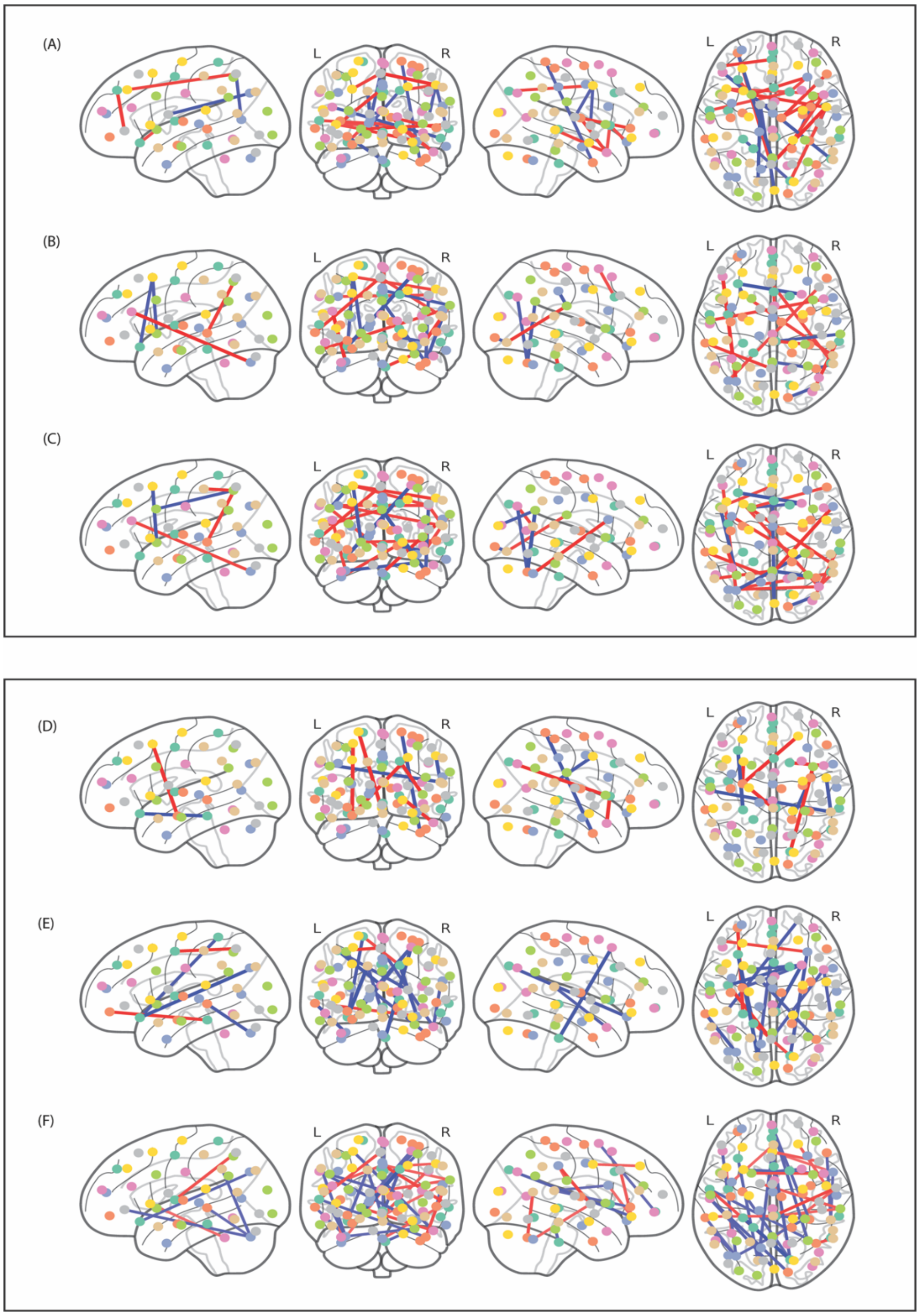

**Supplementary Figure S8. Additional Visualizations for the fMRI Functional Connectivity Features.** Glass brains are shown to depict the functional connectivity features for the (Top Panel) MKL model alone and the (Bottom Panel) difference between the MKL model – SVM model predicting (A, D) the high risk label (CHR-nonconverters + CHR-converters versus control participants); (B, E) CHR-converters versus CHR-nonconverters; and (C, F) CHR-converters versus all others (CHR-nonconverters + control participants).

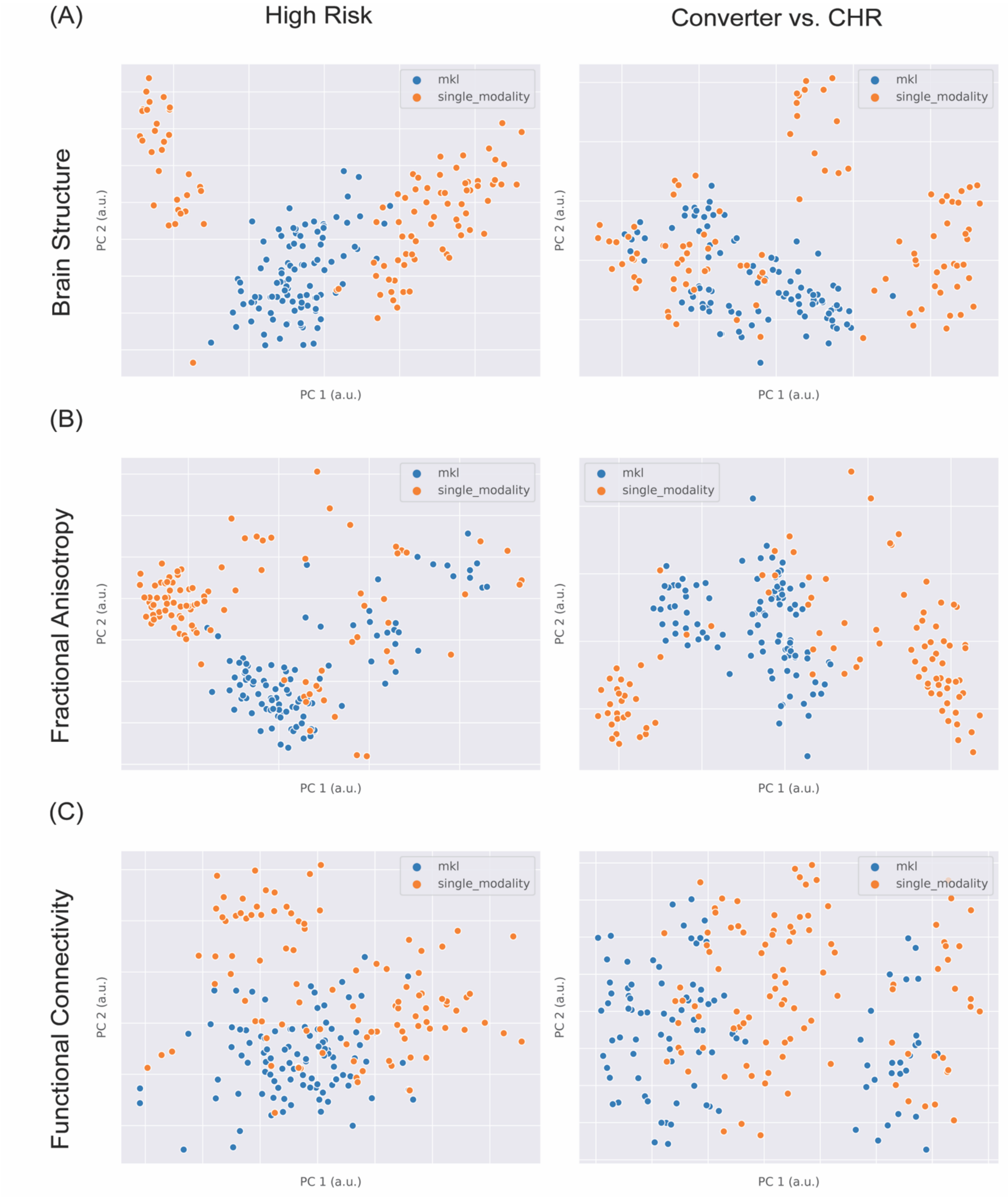

**Supplementary Figure S9. PCA Plots of Weight Vectors Across Training Folds for Unimodal SVM and MKL Models.** Here, they are predicting the high risk label (left column; CHR-nonconverters + CHR-converters versus control participants) and CHR-converters versus CHR-nonconverters (right column) for (A) structural features; (B) DWI features; and (C) fMRI functional connectivity.

#### Supplementary Tables

| CHR LABEL | | | CONVERTER VS. CHR | | | CONVERTER VS. ALL | |
|---|---|---|---|---|---|---|---|
| Left-Thalamus-Proper | -2.28 | | Right-Lateral-Ventricle | 2.74 | | Right-Lateral-Ventricle | 3.45 |
| Left-posteriorcingulate | 2.26 | | Right-medialorbitofrontal | 2.73 | | Right-frontalpole | -2.41 |
| Right-Lateral-Ventricle | 2.23 | | Left-lateraloccipital | -1.96 | | Left-Lateral-Ventricle | 2.36 |
| Left-paracentral | 2.17 | | Left-Lateral-Ventricle | 1.95 | | Right-insula | 2.27 |
| Left-lateralorbitofrontal | -2.06 | | Right-rostralanteriorcingulate | 1.84 | | Left-posteriorcingulate | 2.08 |
| Left-postcentral | 1.94 | | Right-frontalpole | -1.76 | | Right-medialorbitofrontal | 2.07 |
| Right-paracentral | 1.74 | | Right-insula | 1.55 | | Left-lateraloccipital | -1.93 |
| Right-insula | 1.69 | | Right-posteriorcingulate | 1.54 | | Left-parahippocampal | -1.86 |
| Right-Putamen | -1.60 | | Left-Hippocampus | 1.53 | | Left-transversetemporal | 1.77 |
| Left-rostralanteriorcingulate | -1.59 | | Right-inferiortemporal | 1.48 | | Left-insula | 1.68 |
| Right-frontalpole | -1.58 | | Left-posteriorcingulate | 1.47 | | Right-parsorbitalis | -1.50 |
| Left-precentral | 1.51 | | Right-cuneus | -1.43 | | Right-paracentral | 1.43 |
| Right-parsorbitalis | -1.46 | | Left-isthmuscingulate | 1.37 | | Right-rostralanteriorcingulate | 1.41 |
| Right-isthmuscingulate | 1.45 | | Left-inferiortemporal | 1.36 | | Right-inferiortemporal | 1.40 |
| Left-temporalpole | -1.44 | | Left-transversetemporal | 1.33 | | Left-inferiortemporal | 1.39 |

**Supplemental Table S1. Structural features predicting high risk and conversion to psychosis in the MKL model**. The top 15 of 84 feature weights are shown based on descending absolute value for (left to right) regions for each of the analyses predicting (1) the high risk label (CHR-nonconverters + CHR-converters) relative to control participants; (2) the converter label (CHR-converter) relative to other CHR participants (CHR-nonconverters); and (3) the converter label (CHR-converters) vs. all other participants (CHR-nonconverters + controls).

| | Controls v. CHR | | All v. Converters | | CHR: Non-Converters v. Converters | |
|---|---|---|---|---|---|---|
| FEATURE | AUC | p-value | AUC | p-value | AUC | p-value |
| Structural | 0.53 | 0.28 | **0.65** | **0.03** | 0.63 | 0.06 |
| FA | **0.61** | **0.03** | 0.58 | 0.17 | **0.66** | **0.04** |
| MD | 0.41 | 0.96 | 0.48 | 0.61 | 0.56 | 0.25 |
| fMRI | **0.62** | **0.02** | 0.57 | 0.20 | **0.65** | **0.04** |
| MKL(no MD) | **0.66** | **0.01** | **0.68** | **0.01** | **0.73** | **0.01** |
| SVM (no MD) | **0.66** | **0.01** | 0.58 | 0.15 | **0.67** | **0.03** |

**Supplementary Table S2. Comparing Accuracy Across All Analyses.** AUC and significance (p-value) are reported for all individual features (rows) and analyses (columns). *Features: struct = structural features, FA = diffusion fractional anisotropy, MD = diffusion mean diffusivity, fMRI = functional magnetic resonance imaging, MKL = multiple kernel learning model, SVM = support vector machine model.*

| **Controls v. CHR** | | |
|---|---|---|
| | Controls (n=31) | HR (n=43) |
| n Female/Male *ns* | 11 / 20 | 9 / 34 |
| Mean Age (SD)* | 23.23 (3.57) | 21.30 (4.08) |
| Mean Education in Years (SD)* | 4.20 (1.38) | 3.25 (1.60) |
| n Minority/Caucasian† *ns* | 17 / 8 | 25 / 16 |
| **All Participants v. Converters** | | |
| | Non-Converters (n=63) | Converted (n=11) |
| n Female/Male *ns* | 16 / 47 | 4 / 7 |
| Mean Age (SD) *ns* | 22.19 (3.74) | 21.18 (3.67) |
| Mean Education in Years (SD) *ns* | 3.60 (1.56) | 3.70 (1.77) |
| n Minority/Caucasian† *ns* | 35 / 21 | 7 / 3 |
| **CHR: Non-Converters v. Converters** | | |
| | Not Converted (n=32) | Converted (n=11) |
| n Female/Male *ns* | 5 / 27 | 4 / 7 |
| Mean Age (SD) *ns* | 21.66 (5.28) | 21.18 (3.67) |
| Mean Education in Years (SD) *ns* | 3.1 (1.54) | 3.70 (1.77) |
| n Minority/Caucasian† *ns* | 18 / 13 | 7 / 3 |
| Mean SIPS Pos (SD) *ns* | 11.53 (4.94) | 13.55 (5.34) |
| Mean SIPS Neg (SD) *ns* | 14.19 (6.78) | 17.36 (9.77) |
| Mean SIPS Disorganized (SD) *ns* | 7.68 (4.53) | 9.73 (4.47) |
| Mean SIPS General (SD) *ns* | 9.45 (4.14) | 11.55 (5.30) |
| Mean SIPS GAF (SD) *ns* | 47.87 (8.22) | 43.55 (10.23) |

* Denotes significant difference at p<0.05

*ns* denotes not significantly different at p<0.05

*†Minority status is not reported for all participants.*

CHR = clinical high risk; SD = standard deviation; SIPS = Structured Interview for Psychosis-Risk Syndrome

**Supplementary Table S3**. **Demographic and symptoms across groups**. Mean and standard deviation (SD) for demographics and symptoms are reported. Chi-square tests were used to assess gender and minority status. T-tests assessed all other comparisons.
